# Evidence of antigenic imprinting in sequential Sarbecovirus immunization

**DOI:** 10.1101/2020.10.14.339465

**Authors:** Huibin Lv, Ray T. Y. So, Meng Yuan, Hejun Liu, Chang-Chun D. Lee, Garrick K. Yip, Wilson W. Ng, Ian A. Wilson, Malik Peiris, Nicholas C. Wu, Chris K. P. Mok

## Abstract

Antigenic imprinting, which describes the bias of antibody response due to previous immune history, can influence vaccine effectiveness and has been reported in different viruses. Give that COVID-19 vaccine development is currently a major focus of the world, there is a lack of understanding of how background immunity influence antibody response to SARS-CoV-2. This study provides evidence for antigenic imprinting in *Sarbecovirus*, which is the subgenus that SARS-CoV-2 belongs to. Specifically, we sequentially immunized mice with two antigenically distinct *Sarbecovirus* strains, namely SARS-CoV and SARS-CoV-2. We found that the neutralizing antibodies triggered by the sequentially immunization are dominantly against the one that is used for priming. Given that the impact of the background immunity on COVID-19 is still unclear, our results will provide important insights into the pathogenesis of this disease as well as COVID-19 vaccination strategy.

## INTRODUCTION

Coronavirus disease 2019 (COVID-2019) pandemic is an ongoing global public health crisis and has imposed a huge burden on the world economy. The causative agent of COVID-2019, SARS-CoV-2, belongs to the subgenus *Sarbecovirus* of the genus *Betacoronavirus* and has nearly 80% sequence identity with another *Sarbecovirus*, SARS-CoV, which caused a global epidemic in 2003 (Peiris et al., 2003). While many SARS-CoV-2 vaccine candidates are being actively developed (Funk et al., 2020), the influence of immune history on vaccine effectiveness remains elusive. In fact, it is now well recognized that the effectiveness of a given influenza vaccine varies among people with different influenza immunization or infection histories (Henry et al., 2018). Such phenomenon is known as antigenic imprinting, which describes immunological memory induced by primary innate and adaptive immune responses to the first encounter with a microbial pathogen or vaccination that can be retained over a person’s life time. Antigenic imprinting has also been reported in dengue virus (Mongkolsapaya et al., 2003), HIV (Klenerman and Zinkernagel, 1998) and influenza virus (Gouma et al., 2020). Since SARS-CoV-2 may become a seasonal human coronavirus (Kellam and Barclay, 2020) and other zoonotic *Sarbecovirus* strains continue to post a pandemic threat (Menachery et al., 2015), it is important to understand how antigenic imprinting may affect the antibody response to *Sarbecovirus*. In this study, we explored the antigenic imprinting effect of *Sarbecovirus* by characterizing the antibody response from mice that were sequentially immunized two antigenically distinct *Sarbecovirus* strains, namely SARS-CoV and SARS-CoV-2.

## RESULTS

Balb/c mice (6-8 weeks of age) were intraperitoneally (i.p.) immunized twice, with viruses plus adjuvant Addavax (Wu et al., 2019). Four immunization schemes were explored: 1) two rounds of SARS-CoV (SARS-CoV homologous prime-boost); 2) first round with SARS-CoV and second round with SARS-CoV-2 (heterologous SARS-CoV-prime, SARS-CoV-2-boost); 3) two rounds of SARS-CoV-2 (SARS-CoV-2 homologous prime-boost); and 4) first round with SARS-CoV-2 and second round with SARS-CoV (heterologous SARS-CoV-2-prime, SARS-CoV-boost). Plasma samples were collected and antibody immune responses were measured at day 14 after the second round of immunization. Compared to round one of immunization (Lv et al., 2020), the second round homologous boost induced higher homologous binding and neutralizing antibody titers (p <0.05, two tailed t test; Figure S1a-d). These results suggest that SARS-CoV or SARS-CoV-2 specific memory B cells can be recalled and produce neutralizing antibodies during the second round of immunization.

While all our four prime-boost vaccination schemes resulted in cross-reactive RBD-binding antibodies (Figure 1a-d), the virus that was used for priming seemed to dictate the neutralizing antibody response after boosting, regardless of the virus that was used for the boost (Figure 1e-h). For example, no matter whether SARS-CoV or SARS-CoV-2 was used for boosting, the neutralizing antibody response was much stronger against SARS-CoV if SARS-CoV was used for priming (Figure 1e-f), and stronger against SARS-CoV-2 if SARS-CoV-2 was used for priming (Figure 1g-h). Interestingly, while one round of SARS-CoV immunization was sufficient to elicit a detectable SARS-CoV neutralization response (Figure S1c), a SARS-CoV neutralization response was undetectable when SARS-CoV was used as a heterologous boost after priming with SARS-CoV-2 (Figure 1h). A simliar observation was made for SARS-CoV-2 (Figure 1f and S1d).

**Figure. 1.**
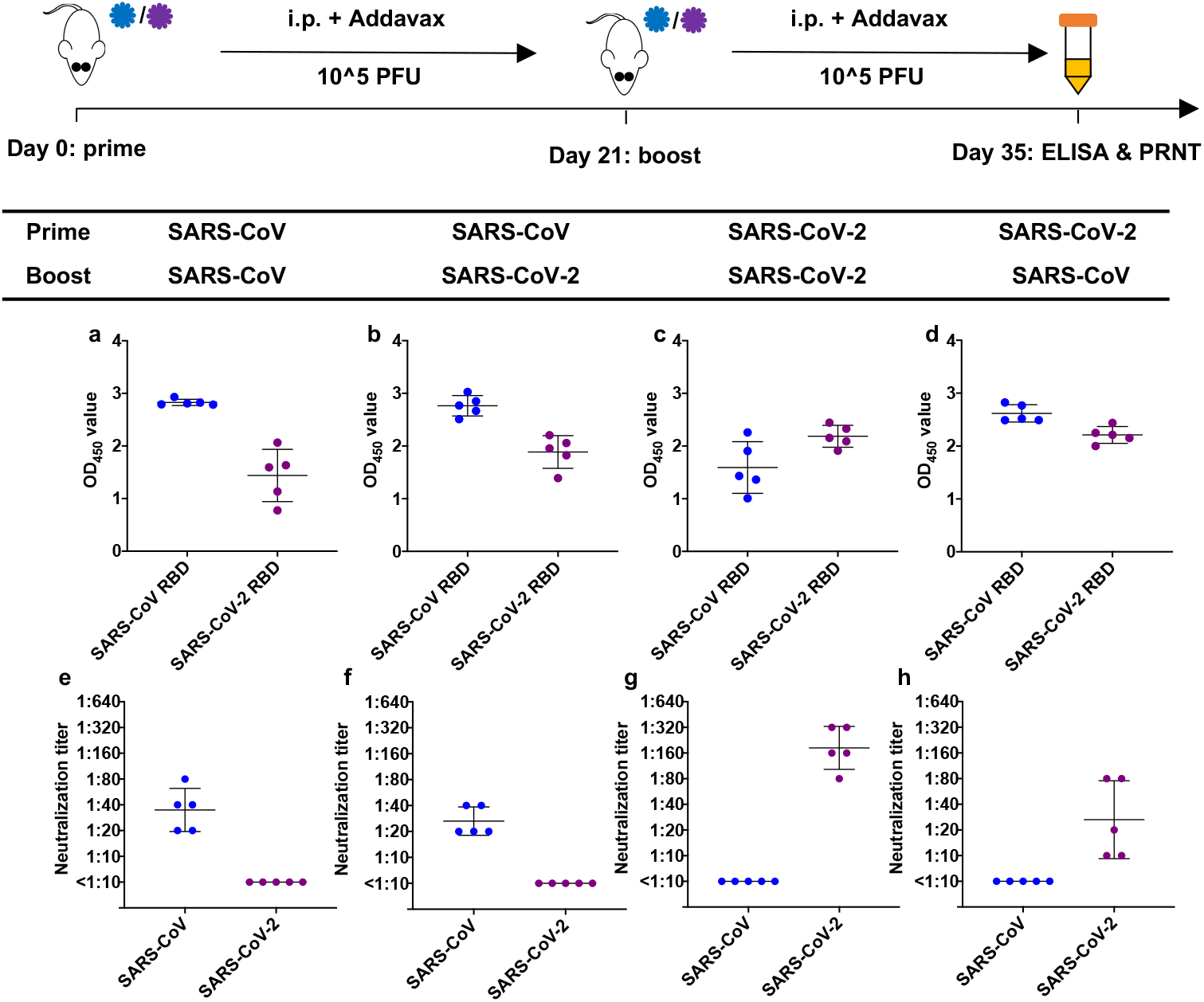
ELISA binding and neutralizing titers of homologous and heterologous sequential immunization with SARS-CoV and SARS-CoV-2. **a-d** RBD (receptor binding domain) proteins from SARS-CoV and SARS-CoV-2 were used as the antigen coating on the ELISA plates. Binding of RBD to 1:100 diluted plasma sample was analyzed from 5 mice immunized using **(a)** SARS-CoV homologous prime-boost, **(b)** heterologous SARS-CoV-prime, SARS-CoV-2-boost, **(c)** SARS-CoV-2 homologous prime-boost, and **(d)** heterlogous SARS-CoV-2-prime, SARS-CoV-boost. The mean OD_450_ value of two replicates are shown. **e-h** Neutralizing titers of plasma samples from mice immunized with **(e)** SARS-CoV homologous prime-boost, **(f)** heterologous SARS-CoV-prime, SARS-CoV-2-boost, **(g)** SARS-CoV-2 homologous prime-boost, and **(h)** heterologous SARS-CoV-2-prime, SARS-CoV-boost, were analyzed by a PRNT (plaque reduction neutralization test) assay. Each data point in the figure represents the mean of two replicates. Error bars represent standard deviation.

Angiotensin-converting enzyme 2 (ACE2) is the host receptor for SARS-CoV and SARS-CoV-2 entry. Previous study has shown that neutralizing activity of sera correlates with the ACE2-competition activity (Tan et al., 2020). ACE2-competition assay was then performed for four groups of plasma samples. Briefly, the binding of plasma samples were tested against RBD and RBD/ACE2 complex. Plasma samples with a stronger ACE2-competition activity should show a greater reduction in binding to RBD/ACE2 complex compared to RBD alone. When binding to SARS-CoV RBD was tested, plasma samples from mice that were primed with SARS-CoV had stronger ACE2-competition activity than did plasma samples from mice that were primed with SARS-CoV-2 (Figure 2a). Similarly, when binding to SARS-CoV-2 RBD was tested, plasma samples from mice that were primed with SARS-CoV-2 had stronger ACE2-competition activity than did plasma samples from mice that were primed with SARS-CoV (Figure 2b). These results indicate that the heterologous boost predominantly induces antibodies to conserved regions outside of the ACE2-binding site that have minimum neutralizing activity. One such example is CR3022, which has strong cross-reactive binding activity to a conserved epitope on RBD, but has weak neutralization activity to SARS-CoV and undetectable neutralization activity to SARS-CoV-2 (Yuan et al., 2020b).

**Figure. 2.**
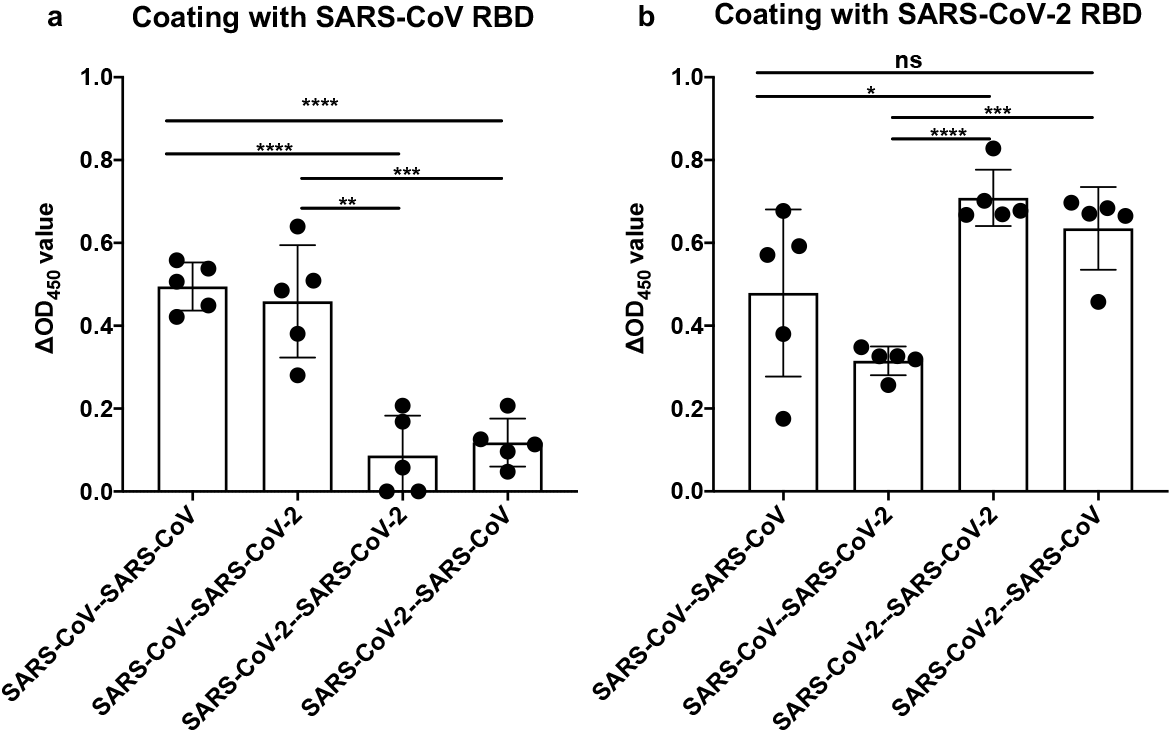
Epitope mapping of the neutralizing and non-neutralizing group. SARS-CoV RBD **(a)** or SARS-CoV-2 RBD **(b)** protein was used as antigen to coat on the 96 well ELISA plate. Four groups of mice plasma samples were added into the plate as primary antibody after with or without 100 ng hACE2 protein blocking. The ΔOD_450_ values were calculated as WT OD_450_ value minus hACE2 blocking OD_450_ value. Each data point in the figure represents the mean of two replicates. P-values were calculated using two-tailed t-test (*P<0.05, **P<0.005, ***P<0.001, ****P<0.0001). Error bars represent standard deviation. Of note, while SARS-CoV-2-SARS-CoV has a stronger ACE2-competition activity to SARS-CoV-2 RBD than SARS-CoV-SARS-CoV, the large standard deviation in SARS-CoV-SARS-CoV makes the difference statistically insigifncant.

Overall, our results suggest that antigenic imprinting can impact the antibody response against *Sarbecovirus*. Specifically, the antibody response to a *Sarbecovirus* strain can be suboptimal if there exists a prior immune history against an antigenically distinct or drifted *Sarbecovirus* strain. This study has important implications for vaccine development against the ongoing COVID-19 pandemic as well as for *Sarbecovirus* and other coronaviruses in general.

## ACKNOWLEDGEMENTS

This work was supported by Calmette and Yersin scholarship from the Pasteur International Network Association (H.L.), International Cooperation and Exchange of the National Natural Science Foundation of China (8181101118), Research Grants Council of the Hong Kong Special Administrative Region, China (Project no. T11-712/19-N) (to J.S.M.P), NIH K99/R00 AI139445 (N.C.W.), and the Bill and Melinda Gates Foundation OPP1170236 (I.A.W.).

## AUTHOR CONTRIBUTION

H.L., N.C.W., and C.K.P.M. conceived and designed the study. N.C.W., M.Y., H.L. and D.C.L. expressed and purified the proteins. H.L., R.T.Y.S., G.K.Y., W.W.N., and C.R.W., performed the experiments. H.L., R.T.Y.S., N.C.W., and C.K.P.M. analyzed the data. H.L., R.T.Y.S., N.C.W., J.S.M.P., I.A.W., and C.K.P.M. wrote the paper, and all authors reviewed and edited the paper.

---

**Figure. S1.**
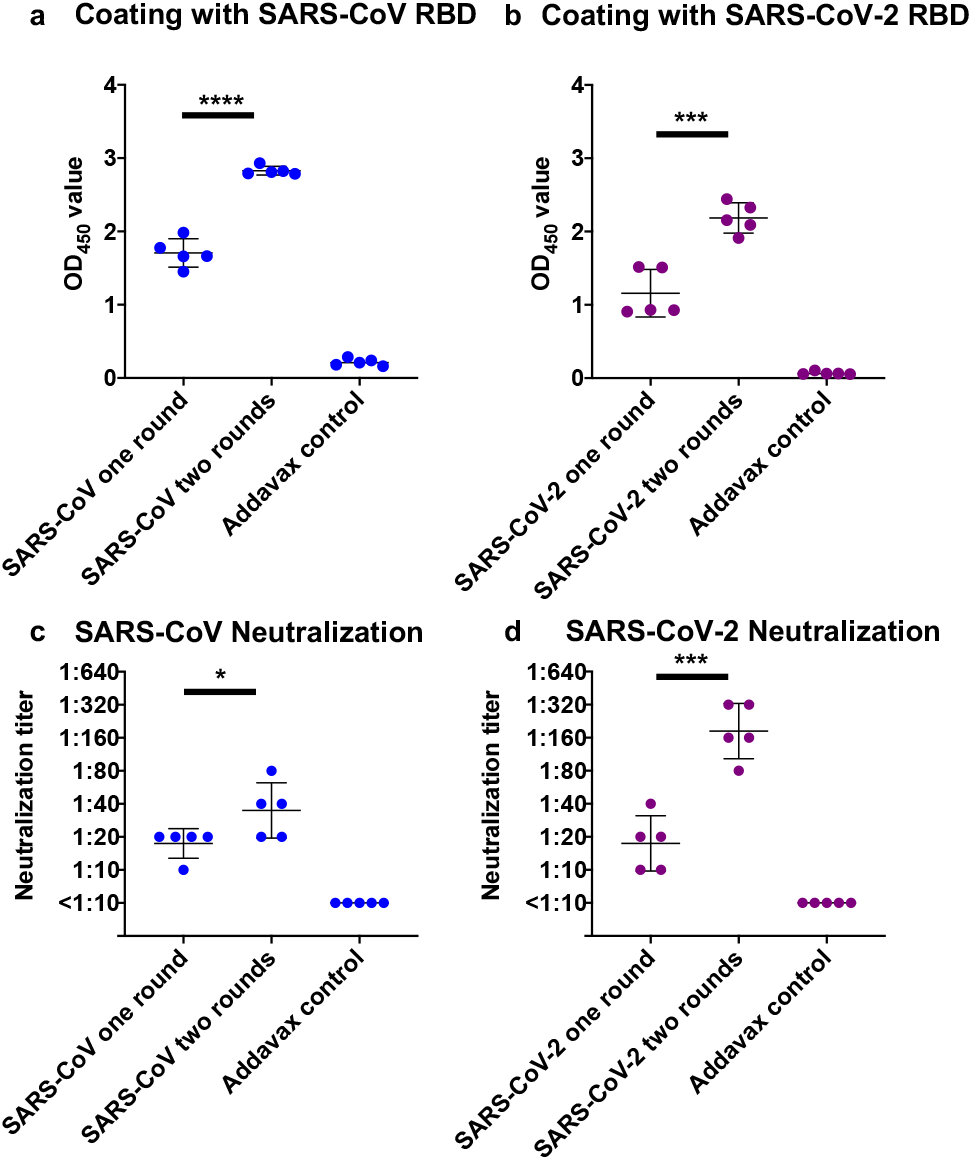
Neutralization titers of one or two rounds of homologous SARS-CoV or SARS-CoV-2 immunization. **a-b** Neutralizing titers of plasma samples from mice immunized by one or two rounds of homologous virus against **(a)** SARS-CoV or **(b)** SARS-CoV-2 were measured by a PRNT assay. Each data point in the figure represents the mean of two replicates. P-values were calculated using two-tailed t-test (*P<0.05, ***P<0.001, ****P<0.0001). Error bars represent standard deviation

## STAR METHODS

### RESOURCE AVAILABILITY

#### Lead Contact

Information and requests for resources and reagents should be directed to and will be fulfilled by the Lead Contact, Chris K. P. Mok.

#### Materials Availability

This study did not generate new unique reagents.

#### Data and Code Availability

NA.

## EXPERIMENTAL MODEL AND SUBJECT DETAILS

### METHOD DETAILS

#### RBD protein expression and purification

The receptor-binding domain (RBD) (residues 319-541) of the SARS-CoV-2 spike (S) protein (GenBank: QHD43416.1) and RBD (residues: 306-527) of the SARS-CoV spike (S) protein (GenBank: ABF65836.1) were cloned into a customized pFastBac vector (Ekiert et al., 2011) and fused with an N-terminal gp67 signal peptide and C-terminal 6×His-tag(Yuan et al., 2020b). A recombinant bacmid DNA was generated using the Bac-to-Bac system (Life Technologies). Baculovirus was generated by transfecting purified bacmid DNA into Sf9 cells using FuGENE HD (Promega), and subsequently used to infect suspension cultures of High Five cells (Life Technologies) at an MOI of 5 to 10. For protein expression, the infected High Five cells were incubated at 28□°C for 72□h with shaking at 110□r.p.m. The supernatant was then concentrated using a 10 kDa MW cutoff Centramate cassette (Pall Corporation). The RBD protein was purified by Ni-NTA, followed by size exclusion chromatography, and buffer exchanged into 20□mM Tris-HCl pH 7.4 and 150□mM NaCl.

#### ACE2 protein expression and purification

The expression of human ACE2 was as previously reported (Yuan et al., 2020a). Briefly, the human ACE2 (residues 19 to 615, GenBank: BAB40370.1) was codon optimized and cloned into phCMV3 vector (Yuan et al., 2020a).The construct was fused with a C-terminal 6xHis tag. The plasmid was transiently transfected into Expi293F cells using ExpiFectamine 293 Reagent (Thermo Fisher Scientific) according to the manufacturer’s manual. At 6 days post-transfection, the supernatant was harvested and then was then washed and eluted with 10 mM and 300 mM Imidazole containing PBS, respectively. The ACE2 eluent was purified by size exclusion chromatography.

#### Mouse immunization

6-8 weeks Balb/c mice were immunized with two rounds 10^5^ pfu of viruses in 150 μL PBS mixing with 50 μL Addavax, including: 1) two rounds of homologous SARS-CoV immunization, 2) two rounds of heterologous immunization with SARS-CoV-prime and SARS-CoV-2-boost, 3) two rounds of homologous SARS-CoV-2 immunization, and 4) two rounds of heterologous immunization with SARS-CoV-2-prime and SARS-CoV-boost, via intraperitoneal (i.p.) route. The plasma samples were collected using heparin tubes on day 35 after the second round of immunization. The experiments were conducted in The University of Hong Kong Biosafety Level 3 (BSL3) facility. This study protocol was carried out in strict accordance with the recommendations and was approved by the Committee on the Use of Live Animals in Teaching and Research of the University of Hong Kong (CULATR 4533-17).

#### ELISA binding assay

ELISA plates (96-well, Nunc MaxiSorp, Thermo Fisher Scientific) were coated overnight with 100 μl of purified recombinant protein in PBS buffer at 1 ng/μl. The plates were then blocked with 100 μl Chonblock buffer (Chondrex Inc, Redmon, US) at room temperature for 1 hours. Each mouse plasma sample was 1:100 diluted in Chonblock buffer, added to the coated ELISA plates, and incubated for 2 hours at 37°C. After three extensive washes with PBS containing 0.1% Tween 20, each well was incubated with the HRP goat anti-mouse secondary antibody (1:5000, Beyotime Biotechnology) for 1 hour at 37°C. The ELISA plates were then washed five times with PBS containing 0.1% Tween 20. Subsequently, 100 μl of TMB buffer (Ncm TMB One; New Cell & Molecular Biotech Co., Ltd) was added into each well. After 15 minutes incubation, 50 μl of 2 M H_2_SO_4_ solution was added to stop the reaction and the plates were analyzed on a Sunrise absorbance microplate reader (Tecan, Ma□nnedorf, Switzerland) at 450 nm wavelength.

#### ACE2-competition ELISA assay

ELISA plates (96-well, Nunc MaxiSorp, Thermo Fisher Scientific) were coated overnight with 100 ng of SARS-CoV or SARS-CoV-2 RBD protein in PBS buffer. The plates were then blocked with 100 μl Chonblock buffer (Chondrex Inc, Redmon, US) at room temperature for 1 hours. After washing, 100 ng of ACE2 protein was added into plate and incubated at 37°C for 2 hours, followed by another 2 hours of 1:200 diluted mouse plasma samples incubation. After three extensive washes with PBS containing 0.1% Tween 20, each well was incubated with the HRP goat anti-mouse secondary antibody (1:5000, Beyotime Biotechnology) for 1 hour at 37°C. The ELISA plates were then washed five times with PBS containing 0.1% Tween 20. Subsequently, 100 μL of TMB buffer (Ncm TMB One; New Cell & Molecular Biotech Co., Ltd) was added into each well. After 15 minutes incubation, 50 μL of 2 M H_2_SO_4_ solution was added to stop the reaction and the plates were analyzed on a Sunrise absorbance microplate reader (Tecan, Ma□nnedorf, Switzerland) at 450 nm wavelength.

#### Plaque reduction neutralization test (PRNT)

Plasma samples were two-fold diluted starting from a 1:10 dilution and mixed with equal volumes of 120 plaque-forming units (pfu) of SARS-CoV-2 as determined by Vero E6 cells. After 1 hour incubation at 37°C, the plasma-virus mixture were added onto Vero E6 monolayers seated in a 24-well cell culture plate and incubated for 1 hour at 37°C with 5% CO_2_. The plasma-virus mixtures were then discarded and infected Vero E6 cells were immediately covered with 1% agarose gel in DMEM medium. After incubation for 3 days at 37°C with 5% CO_2_, the plates were formalin fixed and stained by 0.5% crystal violet solution. Neutralization titers were determined by the highest plasma dilution that resulted in >90% reduction in the number of pfus. The test was performed in a BSL3 facility in the University of Hong Kong.

## REFERENCES

Ekiert, D.C., Friesen, R.H., Bhabha, G., Kwaks, T., Jongeneelen, M., Yu, W., Ophorst, C., Cox, F., Korse, H.J., Brandenburg, B., et al. (2011). A highly conserved neutralizing epitope on group 2 influenza A viruses. Science 333, 843–850.

Funk, C.D., Laferriere, C., and Ardakani, A. (2020). A Snapshot of the Global Race for Vaccines Targeting SARS-CoV-2 and the COVID-19 Pandemic. Front Pharmacol 11, 937.

Gouma, S., Kim, K., Weirick, M.E., Gumina, M.E., Branche, A., Topham, D.J., Martin, E.T., Monto, A.S., Cobey, S., and Hensley, S.E. (2020). Middle-aged individuals may be in a perpetual state of H3N2 influenza virus susceptibility. Nat Commun 11, 4566.

Henry, C., Palm, A.E., Krammer, F., and Wilson, P.C. (2018). From Original Antigenic Sin to the Universal Influenza Virus Vaccine. Trends Immunol 39, 70–79.

Kellam, P., and Barclay, W. (2020). The dynamics of humoral immune responses following SARS-CoV-2 infection and the potential for reinfection. J Gen Virol.

Klenerman, P., and Zinkernagel, R.M. (1998). Original antigenic sin impairs cytotoxic T lymphocyte responses to viruses bearing variant epitopes. Nature 394, 482–485.

Lv, H., Wu, N.C., Tak-Yin Tsang, O., Yuan, M., Perera, R., Leung, W.S., So, R.T.Y., Chun Chan, J.M., Yip, G.K., Hong Chik, T.S., et al. (2020). Cross-reactive antibody response between SARS-CoV-2 and SARS-CoV infections. Cell Rep, 107725.

Menachery, V.D., Yount, B.L., Jr., Debbink, K., Agnihothram, S., Gralinski, L.E., Plante, J.A., Graham, R.L., Scobey, T., Ge, X.Y., Donaldson, E.F., et al. (2015). A SARS-like cluster of circulating bat coronaviruses shows potential for human emergence. Nat Med 21, 1508–1513.

Mongkolsapaya, J., Dejnirattisai, W., Xu, X.N., Vasanawathana, S., Tangthawornchaikul, N., Chairunsri, A., Sawasdivorn, S., Duangchinda, T., Dong, T., Rowland-Jones, S., et al. (2003). Original antigenic sin and apoptosis in the pathogenesis of dengue hemorrhagic fever. Nat Med 9, 921–927.

Peiris, J.S., Lai, S.T., Poon, L.L., Guan, Y., Yam, L.Y., Lim, W., Nicholls, J., Yee, W.K., Yan, W.W., Cheung, M.T., et al. (2003). Coronavirus as a possible cause of severe acute respiratory syndrome. Lancet 361, 1319–1325.

Tan, C.W., Chia, W.N., Qin, X., Liu, P., Chen, M.I., Tiu, C., Hu, Z., Chen, V.C., Young, B.E., Sia, W.R., et al. (2020). A SARS-CoV-2 surrogate virus neutralization test based on antibody-mediated blockage of ACE2-spike protein-protein interaction. Nat Biotechnol 38, 1073–1078.

Wu, N.C., Lv, H., Thompson, A.J., Wu, D.C., Ng, W.W.S., Kadam, R.U., Lin, C.W., Nycholat, C.M., McBride, R., Liang, W., et al. (2019). Preventing an Antigenically Disruptive Mutation in Egg-Based H3N2 Seasonal Influenza Vaccines by Mutational Incompatibility. Cell Host Microbe 25, 836–844 e835.

Yuan, M., Liu, H., Wu, N.C., Lee, C.D., Zhu, X., Zhao, F., Huang, D., Yu, W., Hua, Y., Tien, H., et al. (2020a). Structural basis of a shared antibody response to SARS-CoV-2. Science 369, 1119–1123.

Yuan, M., Wu, N.C., Zhu, X., Lee, C.D., So, R.T.Y., Lv, H., Mok, C.K.P., and Wilson, I.A. (2020b). A highly conserved cryptic epitope in the receptor binding domains of SARS-CoV-2 and SARS-CoV. Science 368, 630–633.

